# Crucial Roles of the Arp2/3 Complex during Mammalian Corticogenesis

**DOI:** 10.1101/026161

**Authors:** Pei-Shan Wang, Fu-Sheng Chou, Fengli Guo, Praveen Suraneni, Sheng Xia, Sree Ramachandran, Rong Li

## Abstract

**HIGHLIGHTS:** Disruption of the Arp2/3 complex impairs cortical development The Arp2/3 complex promotes RGC basal process extension and apical adhesion Loss of Arp2/3 complex leads to altered RGC polarity and cell fate The Arp2/3 complex has non-cell-autonomous and cell-autonomous roles in neuronal migration The Arp2/3 complex enables neuronal cells to migrate on soft or less adhesive substrates

**SUMMARY:** The polarity and organization of radial glial cells (RGCs), which serve as both stem cells and scaffolds for neuronal migration, are crucial for cortical development. However, the cytoskeletal mechanisms that drive radial glial outgrowth and maintain RGC polarity remain poorly understood. Here, we show that the Arp2/3 complex, the unique actin nucleator that produces branched actin networks, plays essential roles in RGC polarity and morphogenesis. Disruption of the Arp2/3 complex in RGCs retards process outgrowth toward the basal surface and impairs apical polarity and adherens junctions. Whereas the former is correlated with abnormal actin-based leading edge, the latter is consistent with blockage in membrane trafficking. These defects result in altered cell fate, disrupted cortical lamination and abnormal angiogenesis. In addition, we present evidence that the Arp2/3 complex is a cell-autonomous regulator of neuronal migration. Our data suggest that Arp2/3-mediated actin assembly may be particularly important for neuronal cell motility in soft or poorly adhesive matrix environment.

## INTRODUCTION

During embryonic neurogenesis, radial glia cells (RGCs), a population of highly polarized stem cells, give rise to most cortical neurons and also serve as their radial migration scaffold (reviewed in (Ayala et al., 2007) and (Rakic, 2003b)). The apically located RGCs maintain their marked apico-basal polarity with a short apical process that forms an end-foot attached to the ventricular zone (VZ) and a long, basal (radial) process that contacts the pia surface, where it is anchored to the basement membrane (Schmechel and Rakic, 1979). The division of RGCs can be either symmetrical, producing two daughter RGCs (to expand the radial glial population), or asymmetrical, producing a RGC and either a daughter neuron (post-mitotic) or an intermediate progenitor cell (IPC, mitotic) (Anthony et al., 2004; Malatesta et al., 2000; Miyata et al., 2001; Noctor et al., 2004; Noctor et al., 2008). The neuron then migrates along the RGC basal process to the cortical plate (CP) and completes the differentiation process. The IPC may undergo multiple rounds of symmetric division to expand in number or to generate neurons (LaMonica et al., 2013). The apical end-feet of RGCs are anchored to each other through adherens junctions (AJs), which is critical for maintaining VZ integrity and RGC identity (Buchman and Tsai, 2007). The basal processes are highly dynamic and are thought to be involved in neuronal positioning (Ayala et al., 2007; Yokota et al., 2009). Abnormalities in RGC polarity would therefore affect both neurogenesis and migration, and may underlie neurodevelopmental disorders and brain tumor (Gotz and Huttner, 2005; Rakic, 2003a; Taylor et al., 2005).

Despite its critical importance in brain development, the molecular mechanisms that regulate RGC polarity, adhesion and basal extension are poorly understood. Recent study indicates that Adenomatous Polyposis Coli (APC) and Cdc42 localize at the tip of RGC basal process and are required for RGC to respond to the polarity maintenance cues, such as neuregulin-1, and to regulate basal end-feet attachment to the pia surface, respectively (Yokota et al., 2010; Yokota et al., 2009). On the ventricular side, the AJ components (cadherens and catenins) and proteins with conserved roles in the regulation of cell polarity and asymmetric cell division (Par3, Par6, aPKC, Cdc42, Numb, etc.) localize to the apical membrane and play critical roles in RGC adhesion and polarity. Loss of these apical polarity proteins resulted in mislocalization or loss of AJ, formation of neuroblastic rosettes, abnormal mitotic entry location, abnormal IPC (intermediate progenitor cell) fate, premature neuronal differentiation and depletion of neural progenitors (Buchman and Tsai, 2007; Bultje et al., 2009; Cappello et al., 2006; Rasin et al., 2007; Zhang et al., 2010).

A likely effector of the above polarity proteins is the actin cytoskeleton, which is also known to play key roles in cell-cell and cell-matrix interactions. A key step in actin polymerization, and thus an important point of regulation *in vivo*, is the nucleation of filaments. An evolutionarily conserved actin nucleator is the Arp2/3 complex, which nucleates branched actin networks. The Arp2/3 complex consists of a stable and stoichiometric assembly of seven polypeptides, including Arp2, Arp3 and Arpc1-5. In the nervous system, the Arp2/3 complex has been shown to be involved in growth cone motility and axon guidance, development of dendritic spines and synapses, and memory decay (Hadziselimovic et al., 2014; Kim et al., 2013; Koch et al., 2014; Korobova and Svitkina, 2008; Lippi et al., 2011; Nakamura et al., 2011; Norris et al., 2009; Rocca et al., 2013; San Miguel-Ruiz and Letourneau, 2014; Spillane et al., 2011; Strasser et al., 2004; Wegner et al., 2008; Yang et al., 2012). The expression level of Arp2/3 complex subunits has been linked to human neurodevelopmental disorder such as Down syndrome (Weitzdoerfer et al., 2002) and brain tumor (Liu et al., 2013). Although the Arp2/3 complex has been studied in varies types of cultured cells, its *in vivo* function in mammalian neurogenesis has not been elucidated due to early embryonic lethality from disruptions of the Arp2/3 complex in mice (Suraneni et al., 2012; Yae et al., 2006). Cdc42 and RhoA, upstream regulators of the Arp2/3 complex, have been shown to control RGC basal process extension and regulate RGC apical adhesion and the cell fate (Cappello et al., 2006; Cappello et al., 2012; Yokota et al., 2010), raising the possibility that the Arp2/3 complex may be crucial for brain development by regulating RGC polarity and morphogenesis.

In this study, we took an approach of conditional gene ablation in mice to dissect the complex function of the Arp2/3 complex during embryonic cortical development. We showed that mouse embryos disrupted of the gene encoding the Arpc2 subunit of the Arp2/3 complex exhibited abnormal corticogenesis. This phenotype is due to defects in RGC apico-basal polarity and radial glial extension, leading to impaired angiogenesis, neurogenesis and neuronal migration. In addition, we showed that the Arp2/3 complex is a cell-autonomous regulatory factor for neuronal migration. We also demonstrated that the Arp2/3 complex plays a role in cellular responsiveness to biochemical and mechanical properties of the environment.

## MATERIALS AND METHODS

### Mice

Mice were cared for according to animal protocols approved by The Stowers Institute for Medical Research. Mice carrying an Arpc2 allele flanked by loxP sites were generated by mating Arpc2^FRT and LoxP^ mice with FLP mice (both obtained from Wellcome Trust Sanger Institute, Hinxton, UK) to delete LacZ and neo between two FRT sites. Arpc2^f/f^Nestin-Cre or Arpc2^f/f^Emx-Cre mouse embryos were generated by mating Arpc2^f/f^ mice with either Arpc2^lox/+^Nestin-Cre^+^ or Arpc2^lox/+^Emx-Cre^+^ mice. Littermate Arpc2^f/f^Nestin-Cre^-^, Arpc2^lox/+^Nestin-Cre^+^ mice are phenotypically normal and are served as control. Arpc2^+/+^Nestin-Cre^+^ mouse embryos are also served as control. Emx1-Cre mice were obtained from The Jackson Laboratory (Bar Harbor, ME).

### Immunohistochemistry, Immunofluorescence and Immunoblot analysis

Mouse embryonic brains were removed and fixed with 4% paraformaldehyde (PFA) in phosphate-buffered saline (PBS) for overnight at 4 °C followed by immunohistochemistry analysis. Neurospheres were plated on poly-D-ornithine and 20 μg/ml laminin coated coverslips, allowed to migrate for 1 hour and then fixed with 4% paraformaldehyde (PFA) in phosphate-buffered saline (PBS) for 15min at room temperature followed by immunofluorescence analysis. Extracts from E13.5 and E15.5 cerebral cortices were prepared in RIPA lysis buffer and were subjected to immunoblot analysis. See Supplemental Data for details.

### *Ex Utero* Electroporation and Preparation of Brain Slice

Approximately 2μl of DNA (2 μg/μl) were injected into the lateral ventricle and electroporated (five 50-ms pulses of 30V at 950-ms intervals). Arp3-GFP plasmid (Plasmid 8462: pEGFP-N1-ACTR3) was obtained from Addgene (Cambridge, MA). Following electroporation, cortices were dissected, coronally sectioned (300μm) in a vibratome (Leica), mounted on Millicell cell culture inserts (Millipore, Bedford, MA), and cultured in MEM/10%FBS for 1 hour followed by Neurobasal medium with 2% B27 for 1 day.

### RGC Process Outgrowth Assay

E14.5 mouse embryonic cortices were dissected and embedded in 100% Matrigel and cultured in Neurobasal medium with 2% B27 for 1-2 days. RGC processes were then imaged by phase-contrast using a Nikon inverted microscope (Nikon ECLIPSE TE2000-E) attached to a live cell incubation chamber. Time-lapse images were processed with ImageJ.

### Transmission Electron Microscopy (TEM)

Mouse brain tissues were harvested and immersion-fixed in 2.5% glutaraldehyde for 2 hours at room temperature. The tissues were washed in PBS and then processed for TEM analysis.

Details about the procedure are provided in Supplemental Data.

### Preparation of GFP-Positive Neurospheres

Mouse cortical neural progenitors were isolated from E14.5 embryonic cortices and cultured as neurospheres. After removal of meninges and cerebellum, cerebral cortex tissue from Arpc2^+/+^Nestin-Cre^+^ and Arpc2^lox/lox^Nestin-Cre^+^ E14.5 mouse embryos were mechanically triturated with a 1-ml Gilson pipette until the cell suspension had no or very few small clumps, filtered through a 70-μm cell strainer, and plated at 5 × 10^4^ cells/ml in a six-well plate (4 ml/well of DMEM/F12 supplemented with 1% B27, 0.5% N2, 20 μg/ml of insulin, 20 ng/ml of basic fibroblast growth factor and 20 ng/ml of epidermal growth factor). After 3 – 4 days, floating neurospheres were passaged at a 1:3 ratio in the same medium. pSicoR (addgene: Plasmid 11579) was used for lentiviral production. Lentiviruses were produced in 293T cells. Neurospheres were dissociated and incubated with lentivirus at 5x10^5^ cells/ml for 6 hours, washed with fresh medium and recovered for 24 hours. After 2 passages, GFP positive cells were sorted by MoFlo Legacy and cultured as neurosphere.

### *In Vitro* Neurosphere Migration Assay

Neurospheres were plated on 0.5-20 μg/ml laminin-coated glass bottom dishes (MatTek Corporation, Ashland, MA), 20 μg/ml laminin-coated elastic surface with a stiffness of 0.2kPa (Softview 35 mm/10 mm glass bottom, 0.2kPa, easy coat, Matrigen, Brea, CA) or 20 μg/ml laminin-coated elastic surface with a stiffness of 1.5, 15 and 28kPa (μ-Dish 35 mm, high, ESS Variety Pack, Ibidi, Madison, WI). Neurospheres were allowed to attach to the bottom and migrate and then imaged by phase-contrast using a Nikon inverted microscope (Nikon ECLIPSE TE2000-E) attached to a live cell incubation chamber during migration. Time-lapse images and kymographs were processed using ImageJ. Neurospheres with similar size (100-150 μm) was selected for quantification. Migration length was measured by selecting the center of mass throughout the length of the movie using the Chemotaxis and Migration tool for ImageJ (Ibidi). Migration length (μm) was then divided by time (min) to obtain migration speed (μm/min). The number of migrating cells was counted by using ImageJ Cell Counter Plugin. Migrating cells are defined as cells that protrude from the boundary of the neurosphere at indicated times.

## RESULTS

### Conditional Ablation of Arpc2 Disrupts Cortical Development

Previous studies demonstrated that conventional gene disruption of the Arpc3 subunit of the Arp2/3 complex results in early embryonic lethality (Suraneni et al., 2012; Yae et al., 2006). We therefore developed a conditional Arp2/3 complex-deficient mouse that allows the function of the complex to be studied at specific developmental stages or tissues. This mouse, purchased originally as a flipper gene-trap line from the Sanger institute (UK), has a floxed allele of Arpc2 subunit, in which Cre-mediated recombination truncates the expression of the protein at amino acid 182 (Fig. S1A). Arpc2 is one of the two central scaffolding subunits of the Arp2/3 complex. Biochemical studies of the Arp2/3 complex in both human and yeast have shown that Arpc2 is essential for the integrity of the entire complex (Goley et al., 2010; Winter et al., 1999). The truncation removes the helix-helix interaction required for the ARPC2/ARPC4 central scaffolds of the complex and mother filament interaction (Daugherty and Goode, 2008; Gournier et al., 2001; Robinson et al., 2001) and is thus predicted to result in complex-complex disruption. To confirm that this truncation results in a null allele, we created the analogous mutation in the budding yeast ARPC2 (Arc35) and confirmed that it produces an Arp2/3 complex-null phenotype (Fig. S1B). Subsequent analysis of the Arpc2^f/f^Nestin-Cre mutant brains confirmed the lack of protein expression and localization of the Arp2/3 complex (see below).

To elucidate the function of the Arp2/3 complex in cortical development, we disrupted Arpc2 by crossing with a Nestin-Cre line (Cre recombinase expression driven by the Nestin enhancer and the human beta globin basal promoter with the 0.3kb intron II) to express the Cre recombinase in the developing RGCs. The Nestin-Cre transgene induced widespread recombination in the CNS neural progenitors from around embryonic day12.5 (E12.5), and loss of Arpc2 was evident in the embryonic cortices of Arpc2^f/f^Nestin-Cre mouse embryos after 13.5 days of gestation (Fig. S2A, S4A). We observed severe intraventricular hemorrhage (IVH) in Arpc2^f/f^Nestin-Cre mouse embryos at E15.5 (Fig. S2B). In addition, thinning of the lateral cortices and enlargement of the lateral ventricles were also apparent from E14.5 (Fig. S2C and S2D). To further verify the roles of the Arp2/3 complex in cortical development, we also disrupted Arpc2 by crossing with an Emx-Cre line, as Emx-Cre expression is more restricted to cortical neural progenitors (De Pietri Tonelli et al., 2008). IVH was again observed in the Arpc2^f/f^Emx-Cre mouse embryos at E14.5 (Fig. S2E). Interestingly, thinning of the lateral cortex and enlargement of the lateral ventricles were not apparent at E14.5 in the Arpc2^f/f^Emx-Cre mouse embryos (Fig. S2E), suggesting that the thinning of the lateral cortices and the enlargement of the lateral ventricles in Arpc2^f/f^Nestin-Cre mouse embryos may be due to pressure generated from severe intraventricular hemorrhage and hydrocephalus.

### Accelerated differentiation of Arpc2-depleted RGCs in association with decreased proliferation and increased apoptosis

To examine the cellular organization of the Arpc2-deficient embryonic cortex, we performed immunostaining of nestin (neural progenitor marker) and Tuj1 (neuronal marker). As neurogenesis first begins, nestin-positive RGCs are confined to the VZ as newly born neurons migrate to the outer layer of the cortex in the control cortex. In contrast, disorganized structures with some ectopic neurogenic rosettes were present in the Arpc2-deficient cortex (Fig. 1A). In addition, there were significant numbers of Tuj1-positive neurons lining the ventricular surface. Arpc2 deletion significantly increased the number of Tuj1-positive neurons at E14.5 (Fig. 1C). Interestingly, whereas E16.5 control cortex exhibited increased Tuj1-positive neurons compared to E14.5 (P<0.01), there was no further increase in Tuj1-positive neurons in the Arpc2-deficient cortex between E14.5 and E16.5. To examine if proliferating neural progenitors were depleted following Arpc2 deletion, we performed immunostaining of Ki67, which is a well-established marker for cells in the cell cycle. Arpc2 deletion significantly decreased the number of Ki67-positive cells at E14.5 (Fig. 1B and 1D). These results suggested premature neuronal differentiation and depletion of neural progenitors following Arpc2 deletion. Since DAPI staining suggested widely occurring cell death in the Arpc2-deficient cortex, we next examined if Arpc2-deficient cortex exhibited increased apoptosis by cleaved caspase-3 immunoreactivity. While apoptotic cells were not detected in the control cortex, there was a significant number of apoptotic cells in the Arpc2-deficient cortex (Fig. S3).

**Figure 1.**
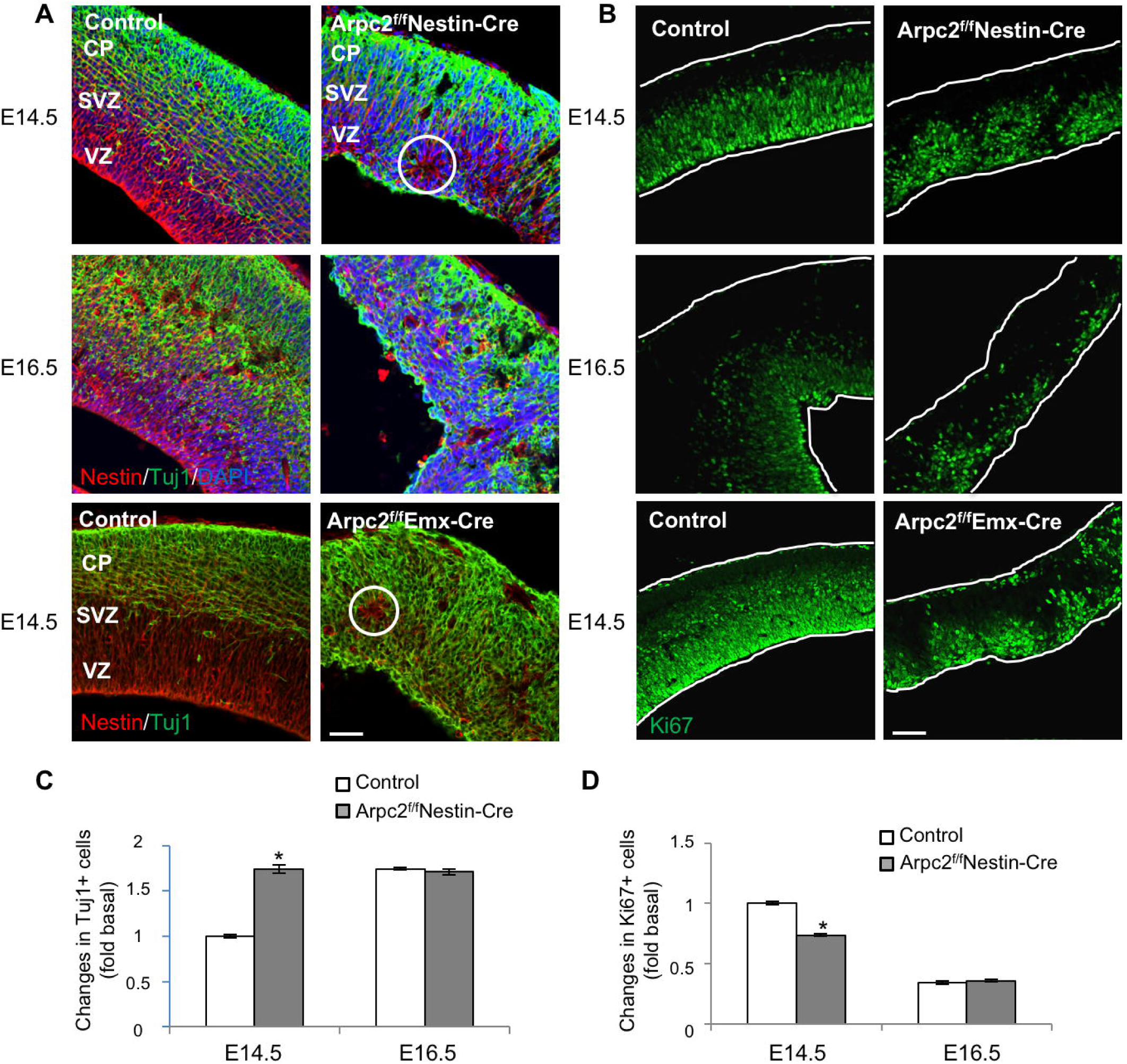
Premature neuronal differentiation and depletion of neural progenitors following Arpc2 deletion. (A) RGCs and cells of the neuronal lineage in E14.5 and E16.5 cortices were labeled with anti-nestin (neural stem cell-specific) and anti-Tuj1 (neuronal lineage-specific) antibodies. In control cortices, a Tuj1-negative ventricular zone (VZ) was clearly shown. In contrast, Arpc2-deficient cortices lack the Tuj1-negative VZ. At E14.5, some ectopic neurogenic rosettes (circle) were observed. SVZ, sub-ventricular zone; CP, cortical plate. (B) Proliferating cells in E14.5 and E16.5 cortices were labeled with anti-Ki67 (proliferation marker) antibody. (C, D) The fluorescence intensity of Tuj1 was measured and the number of Ki67+ cells were counted, followed by normalization to the area of analysis, and shown as fold changes compared to that of the E14.5 control cortex. Note the increase in Tuj1+ and decrease in Ki67+ cells at E14.5 in the Arpc2-deficient cortex. Data shown represent mean ± SEM (n=5); *, significant when compared with control (*p*<0.01, ANOVA test). Scale bar: 50 μm.

### The Arp2/3 complex is required for rapid extension and stability of RGC basal processes

The above phenotypes are consistent with a role for the Arp2/3 complex in RGC morphogenesis. Next, we examined the pattern of localization of the Arp2/3 complex in RGC throughout cortical neurogenesis (E12.5-16.5) by performing immunofluorescent staining of Arp3, a subunit of the Arp2/3 complex. We found that Arp3 was enriched both at the apical and the basal sides of RGCs (Fig. 2A). We also introduced Arp3-GFP into RGC in E14.5 cortex with *ex utero* electroporation to visualize the Arp2/3 complex in individual RGC processes. Localization of Arp3-GFP can be seen throughout the apical and the basal processes as well as cell soma and the nucleus, but it was enriched at both basal and apical end-feet especially at the ventricular surface (Fig. 2B). These observations suggest that the Arp2/3 complex may have multiple roles in the RGCs.

**Figure 2.**
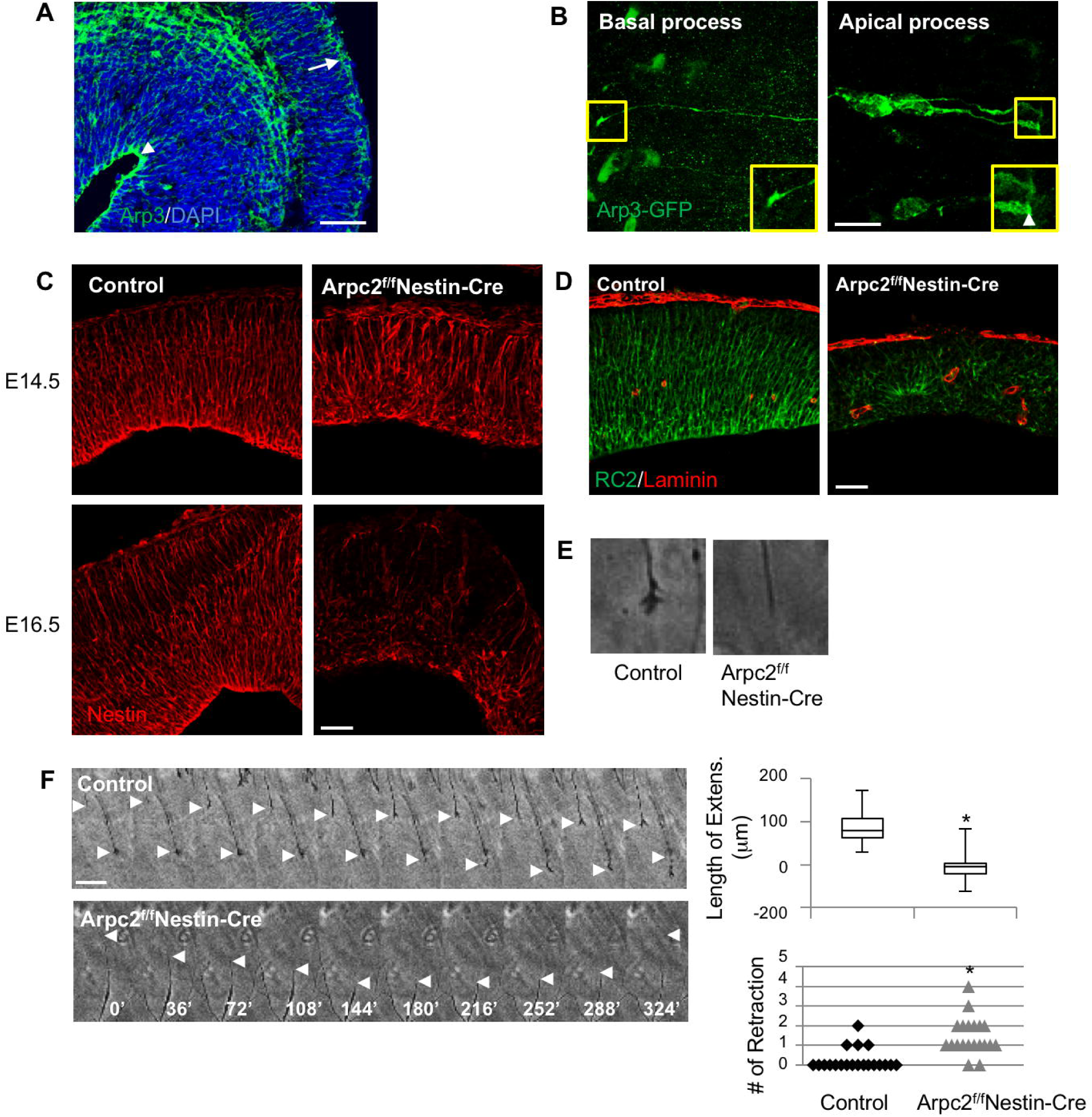
Impaired RGC basal processes following Arpc2 deletion. (A) Immunohistochemical localization of endogenous Arp3 at E16.5 indicated Arp3 expression throughout the developing cerebral wall. Note that Arp3 immunoreactivity was enriched at the apical and basal sides of RGCs (arrowhead, apical end feet; arrow, basal end feet). (B) RGCs in the E14.5 control cortex were electroporated with Arp3-GFP DNA followed by immunostainining with anti-GFP antibody. Arp3 was expressed throughout the RGC basal and apical processes as well as cell soma. It was enriched at the basal end feet (B, left panel) and the apical end feet (B, right panel). (B) RGCs in E14.5 and E16.5 cortices were labeled with anti-nestin antibody. In the control cortex, polarized RGCs span the width of the cerebral wall. In contrast, in the Arpc2-deficient cortex, the polarized organization of RGCs is severely disrupted. At E14.5, many RGCs processes were not extended from the ventricular surface, but from the ectopic rosettes. At E16.5, the majority of the Arpc2-deficient RGC processes were short and misoriented. (D) E14.5 cortices were stained with RC2 (RGC marker) and anti-laminin (pia basement membranes) antibodies. Note the RGC processes in Arpc2-deficient cortex were short and disoriented (many extended from the rosstte). (E, F) *Ex vivo* RGC process outgrowth analysis. Control and Arpc2-deficient cortices were embedded in 100% Matrigel and live imaged. (E) Note that Arpc2-deficient RGC processes were pointy and the dynamic raffles were absent. (F) Montages depicted process outgrowth of control RGCs and Arpc2-deficient RGCs. The length of net extension and the number of retractions in the control and Arpc2-deficient RGC processes within 6 hours of time-lapse recording were measured. Box-plot showing length of net extension of the control RGCs (*n*=20; median 78.8, range 29.8-173.4) and that of the Arpc2-deficient RGCs (*n*=20; median -3.73, range -61.2-82.5). *, significant when compared with controls (*p*<0.01, ANOVA test). Note that shorter length of extension in the Arpc2-deficient RGC processes compared to that of the control RGC processes due to frequent retractions. Scale bar: 50 μm (A), 10 μm (B), 50 μm (C, D) 2.5 μm (E), 25 μm (F).

To further characterize RGC defects resulted from Arpc2 ablation, we used anti-nestin and anti-RC2 antibodies as markers to assess the morphology of the RGC processes. At E14.5, the control developing cortex showed radial organization of the RGC processes that span the width of the cerebral wall, while the Arpc2-deficient cortex exhibited disorganized processes (Fig. 2C). This defect was even further exacerbated at a later stage (E16.5), when the remaining RGCs extended short, misoriented processes and the entire RGC scaffold was drastically abnormal (Fig. 2C). Laminin immunostaining was performed to assess the anchorage of RGCs, labeled by RC2, to the basement membrane (Fig. 2D). Laminin-positive basement membrane, although somewhat discontinuous, was present in the Arpc2-deficient cortex, yet most of the RGC basal processes were not attached to the basement membrane. Therefore, the results suggest that the defective RGC process extension was not due to the lack of basement membrane but more likely due to the lack of normal process extension per se.

To directly examine RGC process extension, control and Arpc2-deficient embryonic cortices were embedded in Matrigel and the *ex vivo* RGC process extension was recorded by time-lapse microscopy. Similar approaches have been used to study glia-guided neuronal migration (Edmondson and Hatten, 1987; Voss et al., 2008; Wichterle et al., 1999). Tip morphologies were apparently different: whereas wild-type RGC basal leading edges were dynamic and exhibited a growth cone-like morphology, most leading processes in mutant RGCs were pointy and lacked dynamic raffles (Fig. 2E and Movie S1-S2). In addition, control RGCs extended their basal processes steadily with an average speed of 0.24±0.1 μm/min (Fig. 2F and Movie S1-S2). By contrast, Arpc2-deficient RGCs extended slightly faster, with average speed of 0.33±0.1 μm/min. However, the final average length of extension of Arpc2-deficient RGC processes was much shorter compared to control RGC processes in 6-hour duration as a result of frequent retraction. These results suggest that the Arp2/3 complex is not only required for the formation of the growth cone-like structure at the tip of the basal process but is also crucial for the stability of the extended processes.

Surprisingly, in addition to an impairment in RGC basal process outgrowth, we also found that the lumen of blood vessels (laminin-positive endothelial cells) in the Arpc2-deficient cortex was enlarged (Fig. S4B) and the pia basement membrane (also labeled by laminin) was broken, suggesting that blood vessel malformation during corticogenesis. To rule out the possibility that blood vessel malformation is due to Cre expression and Arpc2 knock-out in the endothelial cells (which also express nestin), we analyzed the ultrastructure of the blood vessels in Arpc2^f/f^ Emx-Cre cortex. The lumen of the capillary was also enlarged and the capillary walls lined by endothelial cells in the Arpc2^f/f^ Emx-Cre cortex were stretched and thin compared to those in the control cortex (Fig. S4C and S4D). This finding may explain the observed IVH phenotype, and is consistent with a recent report that impaired RGC functions led to abnormal brain angiogenesis and neonatal cerebral hemorrhage (Ma et al., 2013).

### Loss of Arpc2 Results in Disrupted RGC Polarity and Adhesion

The enrichment of the Arp2/3 complex at the apical end-feet of RGCs indicates a possibility that it plays a role in the function of apical end-feet, especially given that the Arp2/3 complex has been shown to be involved in AJ formation and maintenance in cultured epithelial cells (Han et al., 2014; Tang and Brieher, 2012), as well as in the maintenance of PAR asymmetry in *C. elegans* early embryos (Shivas and Skop, 2012). AJ components such as N-cadherin and F-actin, as well as apical polarity proteins such as mPar3, form a continuous apical band in the control cortex (Fig. 3A). In contrast, at E14.5 in the Arpc2-deficient cortex, the apical surface was largely devoid of enrichment for these proteins despite some abnormal accumulation. To further examine whether AJs and the subsequent RGC apical polarity were affected in the Arpc2-deficient cortex, we analyzed the ultrastructural organization of RGC by using thin-sectioning transmission electron microscopy at E14.5. A pseudostratified neuroepithelium was observed in the control cortex (Fig. 3B and 3D). While mitotic cell nuclei migrated to the ventricular surface, elongated interphase cells had their nuclei localized at some distance away from the ventricular surface and maintained connections with neighboring cells by AJs through their apical end-feet. In the Arpc2-deficient cortex, most RGCs lacked AJ or polarized alignment (Fig. 3B and 3C). Interphase, but not mitotic, nuclei were frequently seen to localize at the apical surface. Disorganized AJs and mitotic cells were instead observed to be ectopically located in the rosettes in both VZ and SVZ (Fig. 3D).

**Figure 3.**
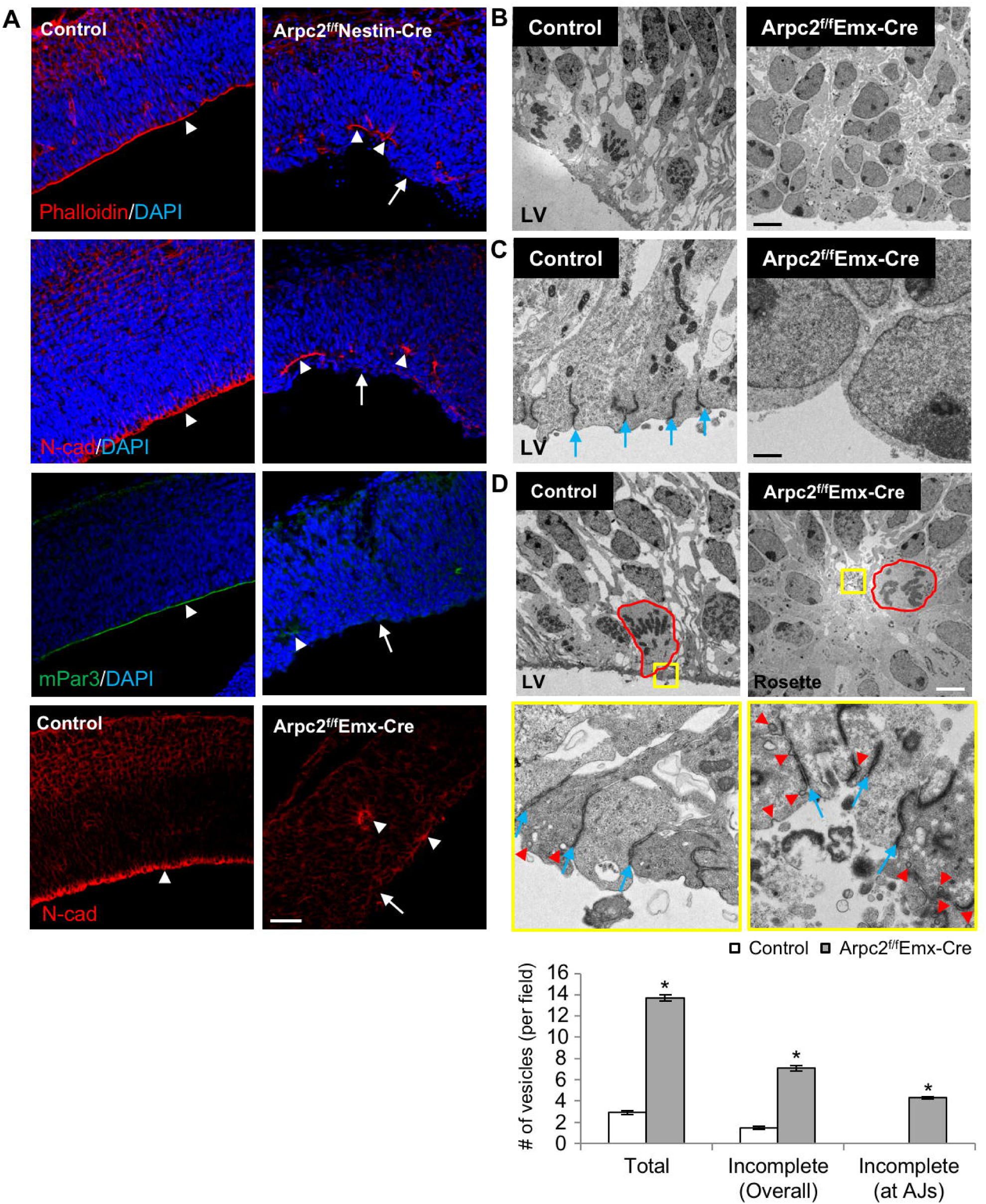
Disrupted RGC apical adhesion following Arpc2 deletion. (A) E14.5 cortices were immunostained with anti-N-cadherin (AJ), anti-mPar3 (apical polarity marker) antibodies or stained with phalloidin (F-actin). Note the loss of actin network, AJ, and apical polarity marker staining as a continuous line but present ectopically after Arpc2 deletion. (B-D) EM analysis of control and Arpc2-deficient VZ at E14.5. (B,C) Control VZ exhibited columnar organization and AJ at the ventricular surface. (D) Arpc2-deficient VZ lacked columnar organization and exhibited neurogenic rosettes. Center of rosettes contained mitotic cells (red outlines) and tangled AJ (middle panel). Note the accumulation of vesicles (red arrowheads) at the adherens junctions and apical surface. Blue arrow, AJ. The size and numbers of vesicles were measured and counted. Values represent mean ± SEM from 10 random fields. *, significant when compared with controls (*p*<0.01, ANOVA test). Note the increase in total vesicles as well as incomplete vesicles at the apical end-feet in the Arpc2-deficient cortex. Also note the significant increase in the incomplete vesicles at the AJs in the Arpc2-deficient cortex. Scale bar: 50 μm (A), 5 μm (B, D), 500 nm (C).

The Arp2/3 complex nucleates the actin filaments for endocytic vesicle scission, a process important for E-cadherin-mediated AJ formation. Indeed, there were more vesicles accumulated in the Arpc2-deficient cortex, near the AJs or the apical membrane of RGC apical end-feet that were ectopically located within the center of rosettes (Fig. 3D). We quantified the number of vesicles at the apical end-feet and showed that endocytic vesicles were increased in the mutant RGCs compared with the control (Fig. 3D). Most of these vesicles remained attached to the plasma membrane, especially at the AJs, in the mutant but not in control RGCs, consistent with a failure in the scission step of endocytosis.

### Loss of Arpc2 Results in Altered Cell Fate and Disorganized Cortical Layers

Since Arp2/3 complex is required for the maintenance of AJs, which have been shown to affect the fate of neural progenitor cells (reviewed in (Kim and Walsh, 2007)), we next analyzed whether the Arpc2-deficient RGCs were able to maintain their stem-cell identity or adopt an IPC fate. As RGCs and IPCs divide at apical and basal positions, respectively, observation of both of these progenitor populations can be accomplished by immunolabeling with anti-phospho-histone H3 (PH3). Arpc2 deletion significantly reduced the number of PH3-positive cells lining the ventricular surface, and increased the number of PH3-positive cells that divide at more basal positions (Fig. 4A, 4B and 4C). Staining of Pax6, an RGC marker, or Tbr2, an IPC marker, showed that Arpc2 deletion leads to a reduction in the number of RGCs (Fig. 4A, 4B and 4D) and a concomitant increase in the number of IPCs whose distribution extends all the way to the ventricular surface (Fig. 4A, 4B and 4E). In fact, most of the dividing cells in the basal area of the Arpc2-deficient cortex were Tbr2-positive IPCs (Fig. 4G). These results suggest that Arp2/3 complex is not required for the division of RGCs or IPCs but is crucial for maintaining the RGC identity. At E16.5, both control and Arpc2-deficient cortices showed decreased numbers of RGCs compared to those at E14.5, and Arpc2 deletion showed a further reduction in the number of RGCs at both time points (Fig. S5A and S5C). The number of IPCs in the Arpc2-deficient cortex increased at E14.5 but decreased at E16.5 (Fig. S5B and S5D). Based on the above-mentioned findings, the reduction in the number of RGCs may be due to exhaustion of RGCs as a result of premature differentiation, decreased proliferation, and/or increased apoptosis.

**Figure 4.**
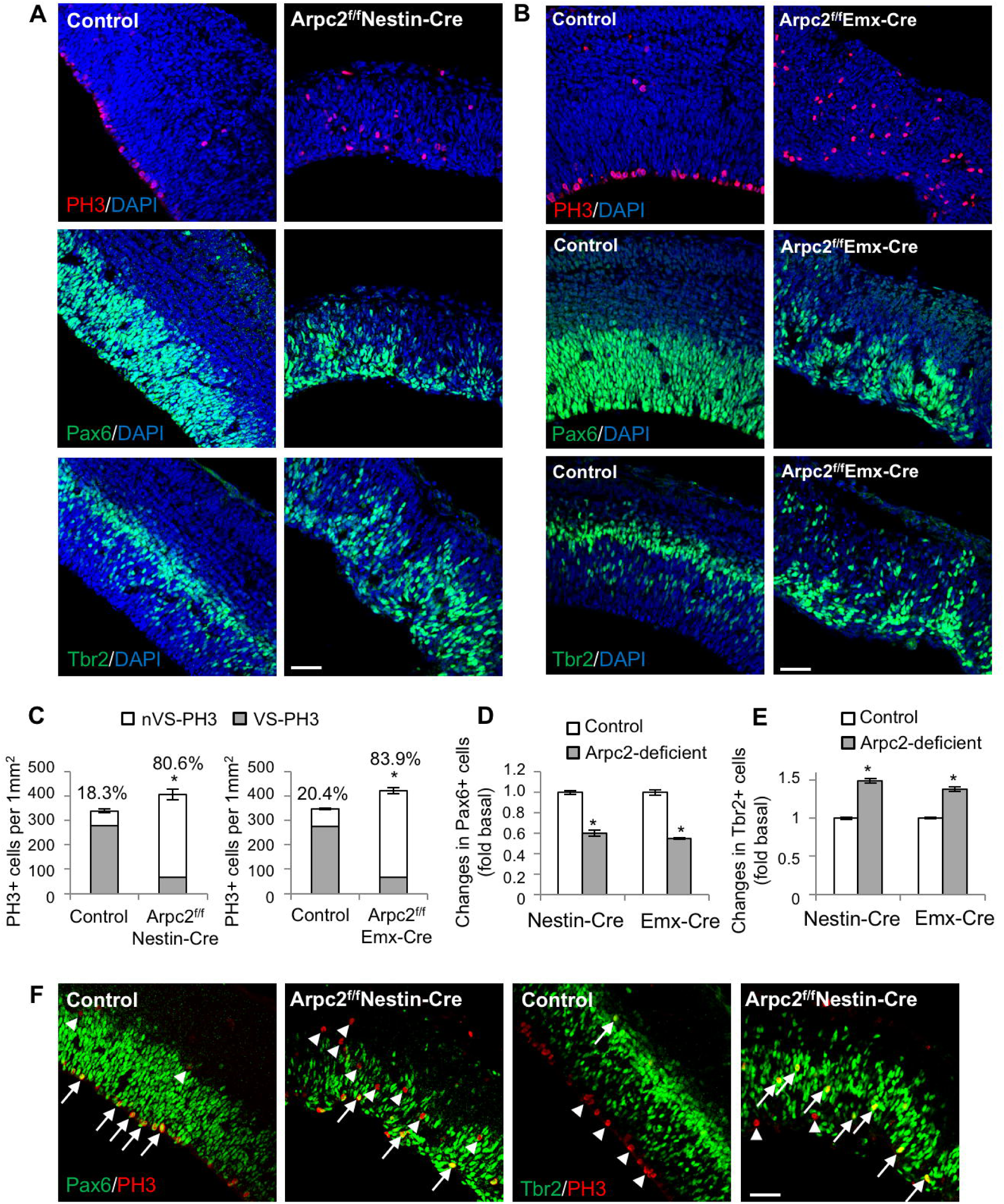
Abnormal mitotic entry location and altered cell fate in the Arpc2-deficient cortex. (A, B) E14.5 cortices were immunostained with anti-phospho H3 (PH3), anti-Pax6 (RGC marker) or Tbr2 (IPC marker) antibodies. (C) The number of PH3+ cells lining the ventricular surface (VS-PH3) or localized at the basal position (nVS-PH3) were counted and normalized to the area of analysis. Note the increased percentage of nVS-PH3+ cells in Arpc2-deficient cortex. (D, E) The number of Pax6+ and Tbr2+ cells were counted and normalized to area of analysis, and showed as fold changes compared to the control cortex. Note the decrease in Pax6+ and the increase in Tbr2+ cells in the Arpc2-deficient cortex. Data shown are mean ± SEM (n=4); *, significant when compared with controls (*p*<0.01, ANOVA test). (G) E14.5 cortices were immunostained with anti-PH3, anti-Pax6 or anti-Tbr2 antibodies. Arrow, PH3 double-positive cells; arrowhead, PH3 single-positive cells. Note the increase in the Tbr2/PH3 double-positive cells in the Arpc2-deficient cortex. Scale bar: 50 μm (A, B, G).

### Loss of Arpc2 results in disrupted cortical lamination and impaired neurogenesis

Disruption in the final neuronal positions was evident in the Arpc2^f/f^Emx-Cre cortex as early as E14.5, even though there was no apparent enlargement in the lateral ventricles in these mutants (Fig. 5A and 5B). To examine cortical lamination following Arpc2 deletion, newly generated cortical neurons were stained with cortical layer-specific markers, Ctip2 and Brn2. In the control cortex, Ctip2-positive and Brn2-positive neurons migrated to distinct laminar positions (Fig. 5C). Both Ctip2-and Brn2-positive neurons were generated in the Arpc2-deficient cortex, but the laminar organization of the neurons in the Arpc2-deficient cortex was severely disrupted. At E16.5, the number of Brn2+ neurons was significantly reduced in the Arpc2-deficient cortex, but the decrease in the number of Ctip2+ neurons in the Arpc2-deficient cortex was not statistically significant (Fig. 5D).

**Figure 5.**
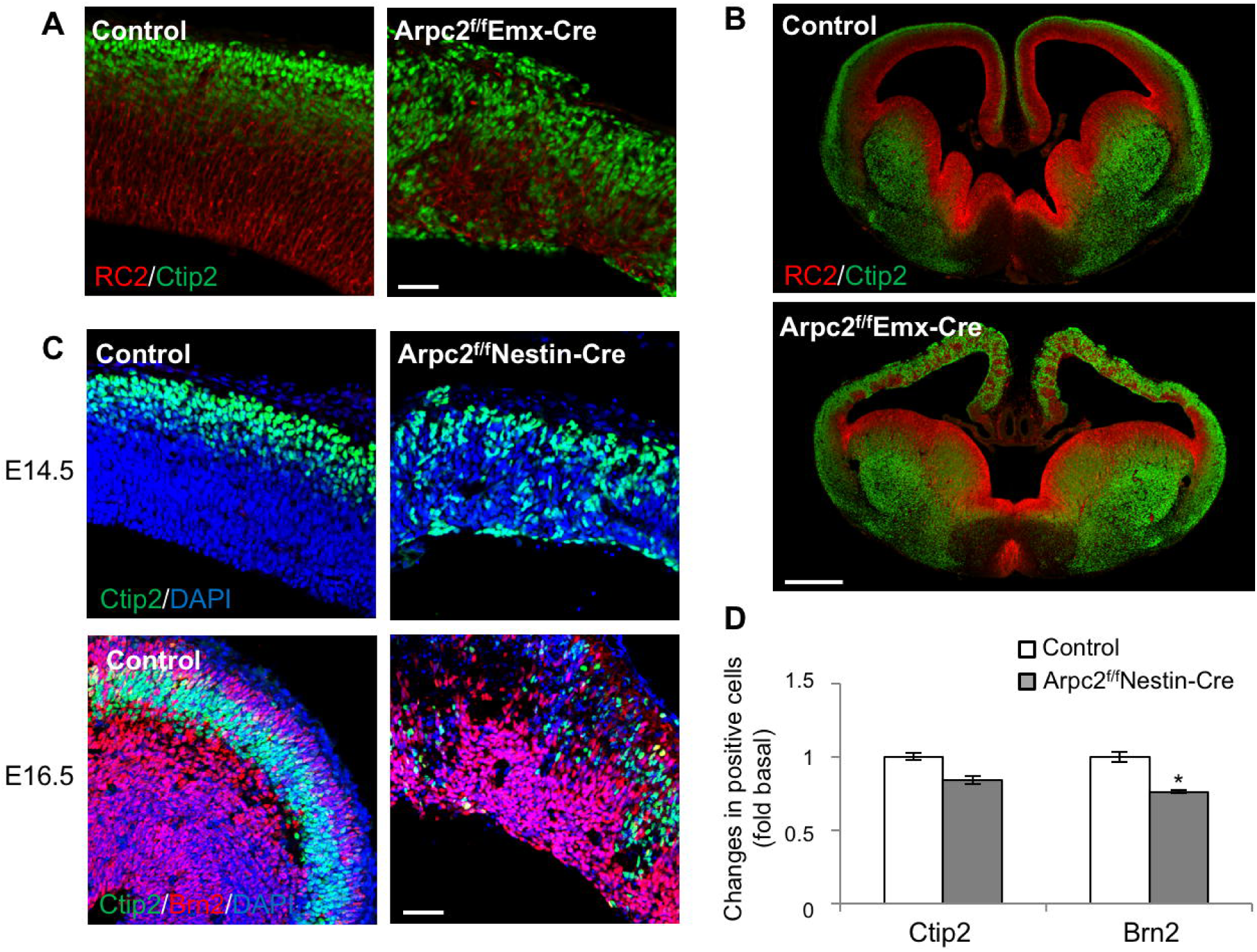
Impaired neurogenesis and disrupted cortical lamination following Arpc2 deletion. (A, B) E14.5 cortices were labeled with RC2 (RGC marker) and anti-Ctip2 (layer V) antibodies. Note the disorganized RGC processes and the cortical neurons in the dorsal region of the Arpc2^f/f^Emx-Cre cortex, but not in the ventral region. (C) E14.5 and E16.5 cortices were immunostained with anti-Ctip2 (layer V), anti-Brn2 (layer II-III) antibodies. Note neurons lining the ventricular surface or throughout the cerebral wall in the mutant cortex, in contrast to the control cortex. Scale bars: 50 μm (A,C), 500 μm (B). (D) The fluorescence intensity of Ctip2 and Brn2 at E16.5 were quantified followed by normalization to the area of analysis, and shown as fold change compared to that of the control cortex. Note the decrease in the Ctip2+ and the Brn2+ cells at E16.5 in the Arpc2-deficient cortex. Data shown are mean ± SEM (*n*=5); *, significant when compared with the controls (*p*<0.05, ANOVA test).

### The Arp2/3 complex has a cell-autonomous role in neuronal migration

The observed neuronal misplacement could be due to disruptions in RGC scaffolding or defects in neuronal motility. The Arp2/3 complex controls actin nucleation at the leading edge of migrating fibroblasts (reviewed in (Bisi et al., 2013; Pollard, 2007; Suraneni et al., 2012; Wu et al., 2012), but the function of the Arp2/3 complex in neuronal migration has not been determined. However, the neuronal migration defect in Arpc2 mutant cortex could simply result from short and disorganized RGC processes rather than reflecting a cell-autonomous role for Arp2/3 complex in the migrating neuronal precursors. To differentiate between these two possibilities, we used an *ex vivo* brain slice culture system with implementation of neurospheres cultured from control and mutant animals (Fig. 6A). The combination allowed us to examine how the Arpc2-deficient neuronal precursors migrate in the wild-type brain environment, which has normal and polarized RGCs. The control neurosphere-derived neuronal precursors were able to migrate toward the cortical plate via two modes: radial migration (RGC dependent) and tangential migration (RGC independent) (Fig. 6B, 6C and Movie S3-S4). Strikingly, the Arpc2-deficient neurosphere-derived neuronal precursors failed to migrate toward the cortical plate. This was not due to a defect in de-adhesion from the sphere, as even cells that exit the sphere, despite being able to extend and retract processes, were unable to migrate (Fig. 6D and Movie S5-S6). Taken together, these results suggest that the Arp2/3 complex is required for the migration of the neuronal precursors along the RGC processes by affecting both the migratory cells and their radial migration tracks formed by the RGC processes.

**Figure 6.**
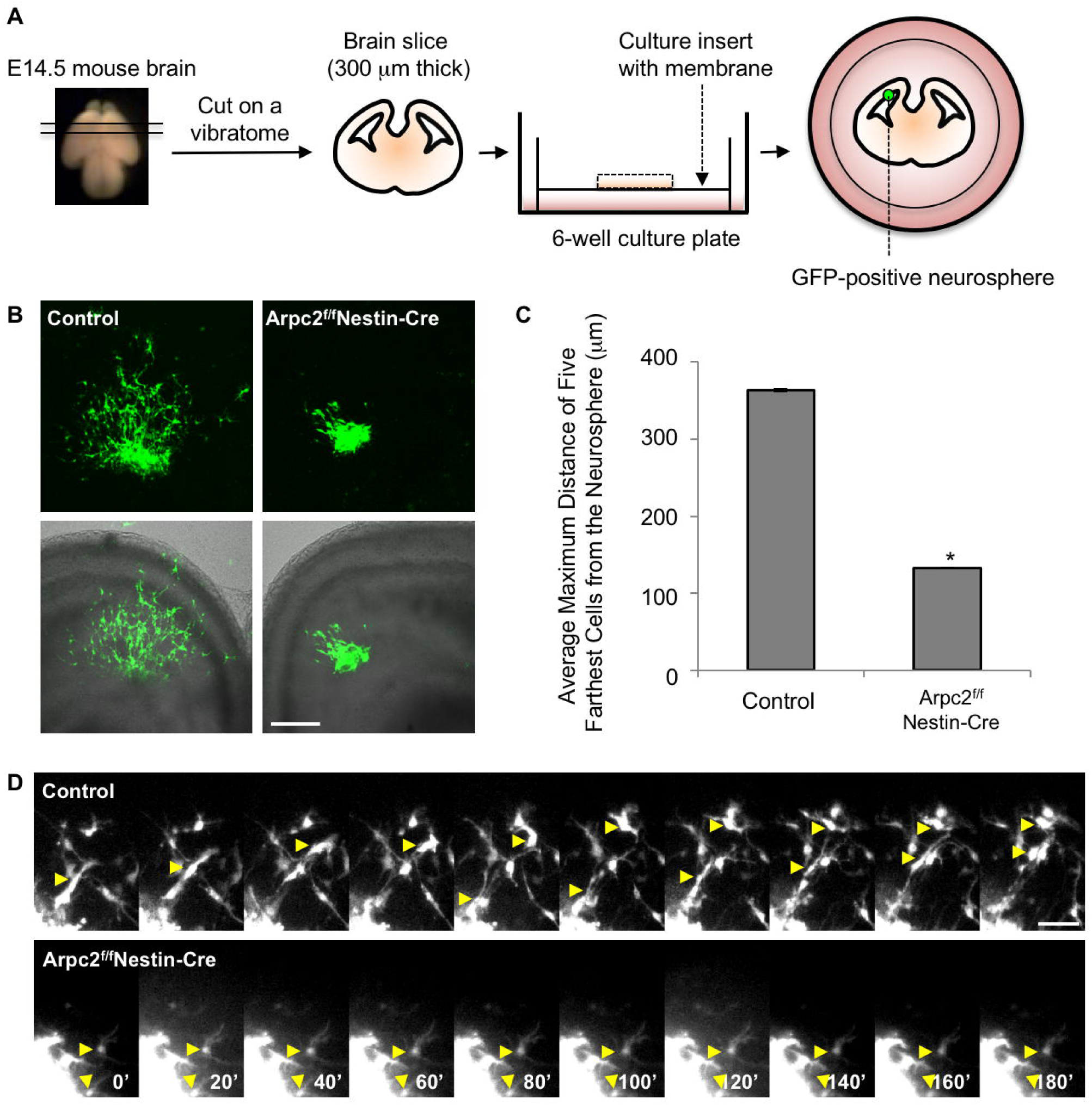
Arpc2-deficient neuronal precursors failed to migrate in the *ex vivo* brain slice culture. (A) A diagram demonstrating the organotypic brain slice co-culture system. E14.5 mouse brain was embedded in 4% agarose and sliced 300μm thick on a vibratome. The brain slices were then cultured on an insert in a 6-well plate. GFP was introduced into the control and the Arpc2-deficient cortical neural progenitors by lentiviral transduction. GFP-positive cells were sorted and cultured as neurospheres and placed on the ventricular surface of the wild-type brain slices. (B) Fluorescence images of migrating neuronal precursors (GFP-positive) after 24 hours of co-culture (upper panel). Combined fluorescence/transmitted light micrograph of brain slice co-cultures (lower panel). Scale bar, 200 μm. (C) Maximum distance of neuronal progenitor migration out of the neurospheres. Values represent mean ± SEM of the five longest migrations from 8 transplanted neurospheres (n=40) from three independent experiments. *, significant compared with the controls (*p*<0.01, ANOVA test). (D) Montages showing migration of two control neuronal precursors (yellow arrowhead) along a RGC basal process, as well as two Arpc2-deficient neuronal precursors (yellow arrowhead) that failed to move. Scale bar: 200 μm (B), 25 μm (D).

### The Arp2/3 complex is particularly crucial for neuronal cells to migrate on soft or less adhesive substrates

To further understand the mechanism by which loss of Arpc2 disrupts neural progenitor cell migration, we established a neurosphere migration assay with phase contrast living imaging to monitor migration of the control and Arpc2-deficient neural progenitor cells on laminin-coated substrate (Fig. 7A). The Arpc2-deficient neural progenitors have the ability to migrate out of the sphere but they moved much more slowly compared to their control neural progenitor counterpart (Fig. 7B-7D and movie S7-S8). Visualization of the Arp2/3 complex by Arpc2 immunostaining showed that the Arp2/3 complex localized in the lamellipodia in the migrating neural progenitors (Fig. 7E). Arpc2-deficient neural progenitors were unable to extend lamellipodia but they were able to generate filopodia-like protrusion. Consistently, the leading edge of Arpc2-deficient neural progenitors was less dynamic (Fig. 7F and movie S9-S10).

**Figure 7.**
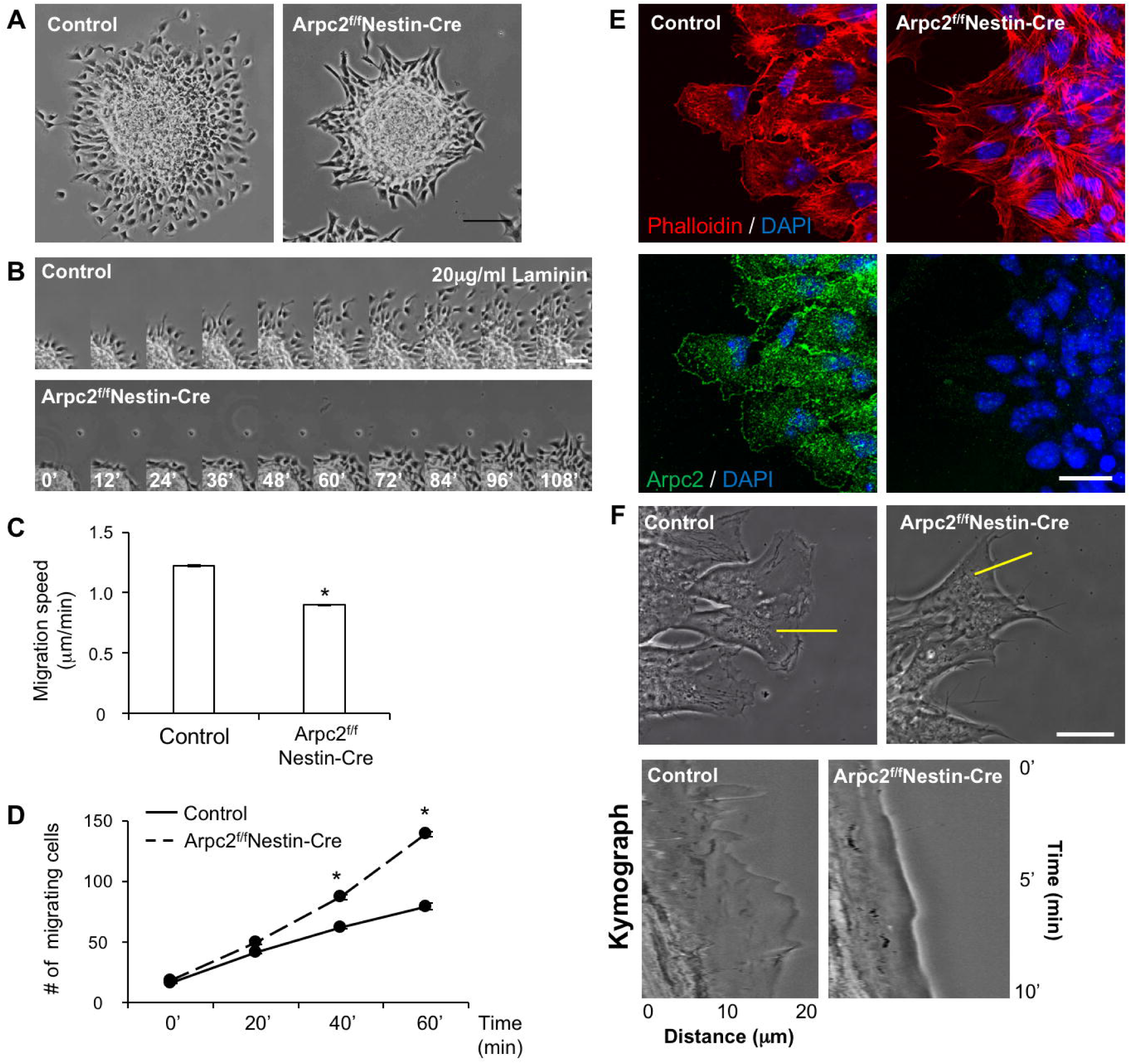
*In vitro* migration of Arpc2-deficient neural progenitors. (A) *In vitro* neurosphere migration assay. Control and Arpc2-deficient neural progenitors migrated on 20 μg/ml laminin-coated glass-bottom dishes for 2 hours. (B) Time-lapse montages of control and Arpc2-deficient neural progenitors migrating on 20 μg/ml laminin-coated glass-bottom dishes. (C) Migrating speed of neural progenitors. Values represent mean ± SEM of the migrating speed of six neural progenitors migrating out of each neurosphere with a total of six neurospheres (*n*=36). Note that Arpc2-deficient neural progenitors migrated slower than the control neural progenitors. (D) Number of neural progenitors migrating out of the neurospheres. Neurospheres of similar size were used. Values represent mean ± SEM of the number of neural progenitors migrating out from six individual neurospheres (*n*=6). (C, D) *, significant compared with controls (*p*<0.01, ANOVA test). (E) Localization of F-actin and endogenous Arpc2 in migrating neural progenitors. Note that Arpc2-deficient neural progenitors are deficient in lamellipodia formation. Also note the localization of Arpc2 in lamellipodia in control but not in Arpc2-deficient neural progenitors. (F) High magnification phase contrast images of control and Arpc2-deficient neural progenitors (top panels). Kymograph analysis along the yellow lines showing local protrusion – retraction cycles (bottom panels). Note the less dynamic leading edge of the mutant cells compared to the control cells. Scale bar: 100 μm (A), 50 μm (B), 20 μm (E,F).

Arpc2-deficient neural progenitors were motile and were able to migrate out of the neurospheres on the laminin-coated glass in this *in* vitro assay but not under the *ex vivo* condition. It is possible that the Arpc2-deficient neural progenitors were unable to respond to environmental cues properly. The physiological concentration of laminin is low in the embryonic brain (Liesi and Silver, 1988), and the brain is one of the softest tissue in the human body (Moore et al., 2010; Spedden et al., 2012). Therefore, we examined whether the Arpc2-deficient neural progenitors failed to migrate in the presence of low laminin concentration (representing low matrix adhesiveness) and of reduced stiffness. Indeed, we found that Arpc2-deficient neural progenitors lost their ability to migrate out of the spheres when they were plated on glass bottom dishes coated with only 0.5 μg/ml, as opposed to 20 μg/ml laminin (Fig. 8A, 8D and movie S11-S12). In addition, Arpc2-deficient neural progenitors also lost their ability to migrate out of the spheres when they were plated on 20 μg/ml laminin-coated elastic surface with a low stiffness index of 0.2 kPa, while they retained the ability to migrate when the stiffness index of the elastic surface was increased to 1.5 kPa (Fig. 8B, 8C, 8D and movie S13-S16). These results suggested a critical role of the Arp2/3 complex in neuronal cell migration in the native brain environment that is both soft and of low laminin concentration.

**Figure 8.**
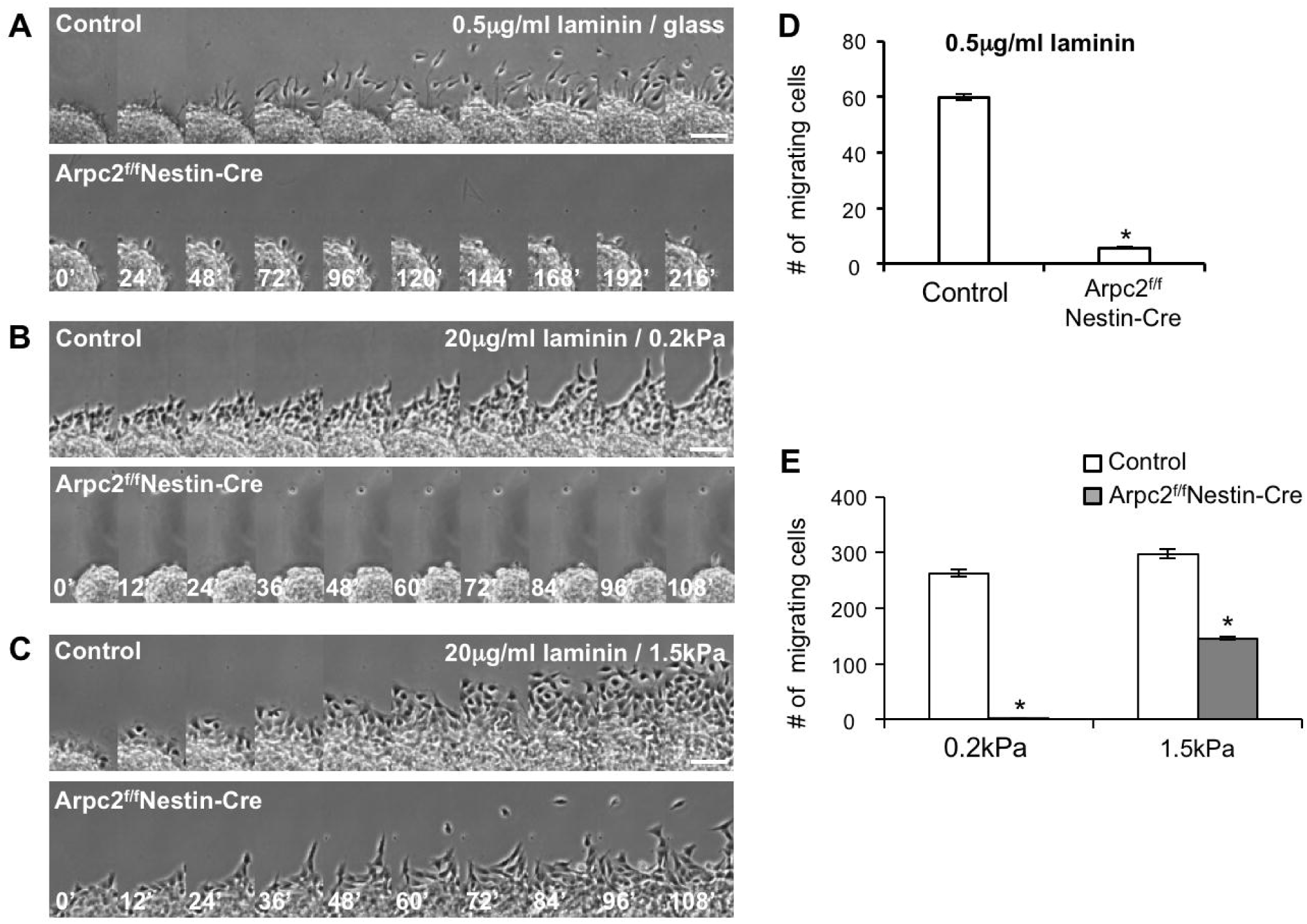
Arpc2-deficient neural progenitors failed to migrate in response to low matrix adhesiveness and low matrix stiffness. Time-lapse montages of control and Arpc2-deficient neural progenitor migration on 0.5 μg/ml laminin-coated glass-bottom dishes (A) and 20 μg/ml laminin-coated elastic surface with a stiffness of 0.2 kPa (B) and 1.5 kPa (C). (D) Number of neural progenitors migrating out of the neurosphere for 6 hours. (E) Number of neural progenitors migrating out of the neurosphere for 2 hours. (D, E) Values represent mean ± SEM of the number of neural progenitors migrating out of the neurospheres (D, *n*=10) (E, *n*=8). *, significant when compared with controls (*p*<0.01, ANOVA test). Note that Arpc2-deficient neural progenitors failed to migrate on low matrix adhesiveness (0.5 μg/ml laminin) or on the soft surface (0.2kPa elastic surface).

## DISCUSSION

Previous studies on the roles of the Arp2/3 complex in neural development have been primarily focused on neuritogenesis, where it was shown that the Arp2/3 complex regulates axon growth cone actin dynamics and guidance, formation of axonal filopodia, and the development of dendritic spines (Korobova and Svitkina, 2008; Nakamura et al., 2011; Norris et al., 2009; Pinyol et al., 2007; San Miguel-Ruiz and Letourneau, 2014; Shakir et al., 2008; Spillane et al., 2011; Strasser et al., 2004; Wegner et al., 2008). Our study, by using conditional Arpc2 gene deletion in mice, has revealed multiple critical roles for the Arp2/3 complex during mammalian corticogenesis. The Arp2/3 complex is required for efficient and stable extension of the RGC basal processes, maintenance of the RGC apico-basal polarity likely through its role in AJ organization, and neuronal migration. Combined defects in these functions result in severely disrupted corticogenesis.

Emerging evidence has suggested that RGC basal process has multiple roles in neurogenesis and neuronal migration (Kosodo and Huttner, 2009). Our data demonstrated that a fast advancing growth cone-like leading edge of RGC requires the Arp2/3 complex and, by inference, formation of a dendritic actin network, suggesting the motility mechanism may be similar to that of neuronal growth cone (Korobova and Svitkina, 2008). It is interesting that the basal extension of Arpc2-deleted RGCs undergo frequent retraction. This phenotype was also observed during neurite outgrowth of neurons isolated from the Arpc2-deficient cortex (P.W. and R.L. unpublished observation). The Arp2/3 complex in some systems was shown to be recruited to microtubules and plays a role in microtubule dynamics (Oelkers et al., 2011; Saedler et al., 2004). Given the facts that both actin and microtubule networks and their interactions are crucial for neurite outgrowth (Gordon-Weeks, 2004), it is possible that Arp2/3 complex-mediated growth cone formation plays a role in stabilizing microtubules in RGC processes.

Another apparent effect of Arpc2 deletion in RGC is the loss of apical AJs and the Par complex, leading to disruptions in neuroepithelial integrity and alterations in progenitor cell fate. This is consistent with the existing evidence that apical polarity and adhesion of RGCs are crucial for RGC self-renewal and the maintenance of the neurogenic niche. For example, it has been shown that deletion of Cdc42, a master regulator of cell polarity, results in retraction of apical processes and an abnormal IPC fate (Cappello et al., 2006). Other apical polarity proteins, such as mPar3, regulate RGC asymmetric division and cell fate via Notch signaling (Bultje et al., 2009). Numb and Numbl inactivation leads to RGC dispersion and disorganized cortical lamination through regulating cadherin recycling (Rasin et al., 2007). Furthermore, N-cadherin-mediated AJs in RGCs regulates ?-catenin signaling and cell fate (Zhang et al., 2010), and loss of cell-cell adhesion or apical polarity leads to a decrease in progenitor cell cycle re-entry (Chenn and Walsh, 2002) and an increase in proneural gene expression (Pierfelice et al., 2011), respectively. Conditional disruption (D6-Cre) of the N-cadherin resulted in the phenotypes similar to our Arpc2 conditional mutant, such as disruption of the AJs localized in the apical end of RGCs, failure of RGCs to extend their bodies or processes between the ventricular zone and the pia surface, as well as scattered mitotic and post-mitotic cells throughout the cortex (Kadowaki et al., 2007). Our observations are also in line with the finding in epithelial cells that WAVE2 and Arp2/3 complex are required for junctional integrity and tension (Han et al., 2014; Verma et al., 2012). In the epithelial cells, Arp2/3 complex is regulated by cortactin and WAVE2, and disruption of these interactions or inhibition of the Arp2/3 complex-mediated actin nucleation abolished actin assembly at the AJs (Han et al., 2014; Tang and Brieher, 2012). Similar mechanisms could underlie the *in vivo* effect of Arpc2 ablation on AJ organization at the ventricular surface.

Arp2/3 complex-mediated actin nucleation is also crucial for endocytosis and exocytosis (Kaksonen et al., 2006; Toret and Drubin, 2006; Zuo et al., 2006). Vesicle trafficking has been shown to be crucial for cortical development and has also been linked to neurodevelopmental disorders (Sheen, 2012). The establishment and the maintenance of AJs require transport of cadherins from the trans-Golgi network to cell-cell contact regions, as well as endocytic recycling of cadherins for continued dynamic remodeling of the AJs in response to tension and cell shape changes (Bryant and Stow, 2004; Lock and Stow, 2005; Paterson et al., 2003). Many of the vesicles accumulated in the Arpc2-deleted RGCs were directly connected to the plasma membrane, consistent with a role for Arp2/3 complex in endocytic vesicle scission. It is possible that Arp2/3 complex maintains RGC apical polarity through regulating endocytic recycling of AJ components. The Arp2/3 complex may also exert its effect on RGC fate by controlling apical trafficking of Delta, a Notch ligand (Rajan et al., 2009) or by controlling asymmetric distribution of PAR complex (Shivas and Skop, 2012).

Arp2/3 complex has been shown to be required for directional migration of fibroblast during chemotaxis and haptotaxis (Suraneni et al., 2015; Suraneni et al., 2012; Wu et al., 2012) and for directional migration of oligodendrocyte progenitor cells in response to the electric field (Li et al., 2015). However, the role of the Arp2/3 complex in the migratory neurons and neural progenitors has not been investigated. Here we demonstrate that neurosphere-derived neuronal precursors and neural progenitors lost their ability to migrate in the *ex vivo* brain slice culture. Our *in vitro* study suggests that Arpc2-deficient neuronal precursors and neural progenitors are intrinsically motile, similar to the Arpc3-deficient fibroblasts and the Arpc2-deficient oligodendrocyte progenitors. However, they failed to migrate in the environment with low matrix adhesiveness and stiffness, which is close to the environment of the embryonic brain used for the *ex vivo* brain slice migration assay. This finding also implies that agents disrupting the Arp2/3 complex-based actin nucleation system may be particularly useful for impeding the precocious migratory ability of glioma cells in the tissue environment of the brain.

## ACKNOWLEDGMENTS

We thank Paco Cambronero and Robb Krumlauf for generating and providing the Nestin-Cre line. This research was supported by NIH grant PO1 GM066311. We also thank Dr. Helen Christou at Harvard Medical School for her kindness in providing lab space, supplies, and reagents for completion of some of the experiments during the paper revision.

## AUTHOR CONTRIBUTIONS

P.-S.W. designed and did most of the experiments and wrote the manuscript. F.-S.C. did the immunoblot analysis, prepared GFP-positive neurospheres, did the *ex vivo* brain slice migration analysis together with P.-S.W, and helped with the manuscript preparation and revision. F.G. did the transmission electron microscopy work. P.S. and S.X. generated the Arpc2^loxp/+^ mouse line. S.R. did the yeast work. R.L. directed the project and wrote the manuscript together with P.-S.W.

## SUPPLEMENTAL MATERIAL

The Supplemental Data include Supplemental Figures, Movies and Materials and Methods.

## REFERENCES

Anthony, T. E., Klein, C., Fishell, G. and Heintz, N. (2004). Radial glia serve as neuronal progenitors in all regions of the central nervous system. Neuron 41, 881–890.

Ayala, R., Shu, T. and Tsai, L. H. (2007). Trekking across the brain: the journey of neuronal migration. Cell 128, 29–43.

Bisi, S., Disanza, A., Malinverno, C., Frittoli, E., Palamidessi, A. and Scita, G. (2013). Membrane and actin dynamics interplay at lamellipodia leading edge. Current opinion in cell biology 25, 565–573.

Bryant, D. M. and Stow, J. L. (2004). The ins and outs of E-cadherin trafficking. Trends in cell biology 14, 427–434.

Buchman, J. J. and Tsai, L. H. (2007). Spindle regulation in neural precursors of flies and mammals. Nature reviews. Neuroscience 8, 89–100.

Bultje, R. S., Castaneda-Castellanos, D. R., Jan, L. Y., Jan, Y. N., Kriegstein, A. R. and Shi, S. H. (2009). Mammalian Par3 regulates progenitor cell asymmetric division via notch signaling in the developing neocortex. Neuron 63, 189–202.

Cappello, S., Attardo, A., Wu, X., Iwasato, T., Itohara, S., Wilsch-Brauninger, M., Eilken, H. M., Rieger, M. A., Schroeder, T. T., Huttner, W. B., et al. (2006). The Rho-GTPase cdc42 regulates neural progenitor fate at the apical surface. Nature neuroscience 9, 1099–1107.

Cappello, S., Bohringer, C. R., Bergami, M., Conzelmann, K. K., Ghanem, A., Tomassy, G. S., Arlotta, P., Mainardi, M., Allegra, M., Caleo, M., et al. (2012). A radial glia-specific role of RhoA in double cortex formation. Neuron 73, 911–924.

Chenn, A. and Walsh, C. A. (2002). Regulation of cerebral cortical size by control of cell cycle exit in neural precursors. Science 297, 365–369.

Daugherty, K. M. and Goode, B. L. (2008). Functional surfaces on the p35/ARPC2 subunit of Arp2/3 complex required for cell growth, actin nucleation, and endocytosis. The Journal of biological chemistry 283, 16950–16959.

De Pietri Tonelli, D., Pulvers, J. N., Haffner, C., Murchison, E. P., Hannon, G. J. and Huttner, W. B. (2008). miRNAs are essential for survival and differentiation of newborn neurons but not for expansion of neural progenitors during early neurogenesis in the mouse embryonic neocortex. Development 135, 3911–3921.

Edmondson, J. C. and Hatten, M. E. (1987). Glial-guided granule neuron migration in vitro: a high-resolution time-lapse video microscopic study. The Journal of neuroscience: the official journal of the Society for Neuroscience 7, 1928–1934.

Goley, E. D., Rammohan, A., Znameroski, E. A., Firat-Karalar, E. N., Sept, D. and Welch, M. D. (2010). An actin-filament-binding interface on the Arp2/3 complex is critical for nucleation and branch stability. Proceedings of the National Academy of Sciences of the United States of America 107, 8159–8164.

Gordon-Weeks, P. R. (2004). Microtubules and growth cone function. Journal of neurobiology 58, 70–83.

Gotz, M. and Huttner, W. B. (2005). The cell biology of neurogenesis. Nature reviews. Molecular cell biology 6, 777–788.

Gournier, H., Goley, E. D., Niederstrasser, H., Trinh, T. and Welch, M. D. (2001). Reconstitution of human Arp2/3 complex reveals critical roles of individual subunits in complex structure and activity. Molecular cell 8, 1041–1052.

Hadziselimovic, N., Vukojevic, V., Peter, F., Milnik, A., Fastenrath, M., Fenyves, B. G., Hieber, P., Demougin, P., Vogler, C., de Quervain, D. J., et al. (2014). Forgetting is regulated via Musashi-mediated translational control of the Arp2/3 complex. Cell 156, 1153–1166.

Han, S. P., Gambin, Y., Gomez, G. A., Verma, S., Giles, N., Michael, M., Wu, S. K., Guo, Z., Johnston, W., Sierecki, E., et al. (2014). Cortactin scaffolds Arp2/3 and WAVE2 at the epithelial zonula adherens. The Journal of biological chemistry 289, 7764–7775.

Kadowaki, M., Nakamura, S., Machon, O., Krauss, S., Radice, G. L. and Takeichi, M. (2007). N-cadherin mediates cortical organization in the mouse brain. Dev Biol 304, 22–33.

Kaksonen, M., Toret, C. P. and Drubin, D. G. (2006). Harnessing actin dynamics for clathrin-mediated endocytosis. Nature reviews. Molecular cell biology 7, 404–414.

Kim, I. H., Racz, B., Wang, H., Burianek, L., Weinberg, R., Yasuda, R., Wetsel, W. C. and Soderling, S. H. (2013). Disruption of Arp2/3 results in asymmetric structural plasticity of dendritic spines and progressive synaptic and behavioral abnormalities. The Journal of neuroscience: the official journal of the Society for Neuroscience 33, 6081–6092.

Kim, S. and Walsh, C. A. (2007). Numb, neurogenesis and epithelial polarity. Nature neuroscience 10, 812–813.

Koch, N., Kobler, O., Thomas, U., Qualmann, B. and Kessels, M. M. (2014). Terminal axonal arborization and synaptic bouton formation critically rely on abp1 and the arp2/3 complex. PloS one 9, e97692.

Korobova, F. and Svitkina, T. (2008). Arp2/3 complex is important for filopodia formation, growth cone motility, and neuritogenesis in neuronal cells. Molecular biology of the cell 19, 1561–1574.

Kosodo, Y. and Huttner, W. B. (2009). Basal process and cell divisions of neural progenitors in the developing brain. Development, growth & differentiation 51, 251–261.

LaMonica, B. E., Lui, J. H., Hansen, D. V. and Kriegstein, A. R. (2013). Mitotic spindle orientation predicts outer radial glial cell generation in human neocortex. Nature communications 4, 1665.

Li, Y., Wang, P. S., Lucas, G., Li, R. and Yao, L. (2015). ARP2/3 complex is required for directional migration of neural stem cell-derived oligodendrocyte precursors in electric fields. Stem Cell Res Ther 6, 41.

Liesi, P. and Silver, J. (1988). Is astrocyte laminin involved in axon guidance in the mammalian CNS? Dev Biol 130, 774–785.

Lippi, G., Steinert, J. R., Marczylo, E. L., D’Oro, S., Fiore, R., Forsythe, I. D., Schratt, G., Zoli, M., Nicotera, P. and Young, K. W. (2011). Targeting of the Arpc3 actin nucleation factor by miR-29a/b regulates dendritic spine morphology. The Journal of cell biology 194, 889–904.

Liu, Z., Yang, X., Chen, C., Liu, B., Ren, B., Wang, L., Zhao, K., Yu, S. and Ming, H. (2013). Expression of the Arp2/3 complex in human gliomas and its role in the migration and invasion of glioma cells. Oncology reports 30, 2127–2136.

Lock, J. G. and Stow, J. L. (2005). Rab11 in recycling endosomes regulates the sorting and basolateral transport of E-cadherin. Molecular biology of the cell 16, 1744–1755.

Ma, S., Kwon, H. J., Johng, H., Zang, K. and Huang, Z. (2013). Radial glial neural progenitors regulate nascent brain vascular network stabilization via inhibition of Wnt signaling. PLoS biology 11, e1001469.

Malatesta, P., Hartfuss, E. and Gotz, M. (2000). Isolation of radial glial cells by fluorescent-activated cell sorting reveals a neuronal lineage. Development 127, 5253–5263.

Miyata, T., Kawaguchi, A., Okano, H. and Ogawa, M. (2001). Asymmetric inheritance of radial glial fibers by cortical neurons. Neuron 31, 727–741.

Moore, S. W., Roca-Cusachs, P. and Sheetz, M. P. (2010). Stretchy proteins on stretchy substrates: the important elements of integrin-mediated rigidity sensing. Developmental cell 19, 194–206.

Nakamura, Y., Wood, C. L., Patton, A. P., Jaafari, N., Henley, J. M., Mellor, J. R. and Hanley, J. G. (2011). PICK1 inhibition of the Arp2/3 complex controls dendritic spine size and synaptic plasticity. The EMBO journal 30, 719–730.

Noctor, S. C., Martinez-Cerdeno, V., Ivic, L. and Kriegstein, A. R. (2004). Cortical neurons arise in symmetric and asymmetric division zones and migrate through specific phases. Nature neuroscience 7, 136–144.

Noctor, S. C., Martinez-Cerdeno, V. and Kriegstein, A. R. (2008). Distinct behaviors of neural stem and progenitor cells underlie cortical neurogenesis. The Journal of comparative neurology 508, 28–44.

Norris, A. D., Dyer, J. O. and Lundquist, E. A. (2009). The Arp2/3 complex, UNC-115/abLIM, and UNC-34/Enabled regulate axon guidance and growth cone filopodia formation in Caenorhabditis elegans. Neural development 4, 38.

Oelkers, J. M., Vinzenz, M., Nemethova, M., Jacob, S., Lai, F. P., Block, J., Szczodrak, M., Kerkhoff, E., Backert, S., Schluter, K., et al. (2011). Microtubules as platforms for assaying actin polymerization in vivo. PloS one 6, e19931.

Paterson, A. D., Parton, R. G., Ferguson, C., Stow, J. L. and Yap, A. S. (2003). Characterization of E-cadherin endocytosis in isolated MCF-7 and chinese hamster ovary cells: the initial fate of unbound E-cadherin. The Journal of biological chemistry 278, 21050–21057.

Pierfelice, T., Alberi, L. and Gaiano, N. (2011). Notch in the vertebrate nervous system: an old dog with new tricks. Neuron 69, 840–855.

Pinyol, R., Haeckel, A., Ritter, A., Qualmann, B. and Kessels, M. M. (2007). Regulation of N-WASP and the Arp2/3 complex by Abp1 controls neuronal morphology. PloS one 2, e400.

Pollard, T. D. (2007). Regulation of actin filament assembly by Arp2/3 complex and formins. Annual review of biophysics and biomolecular structure 36, 451–477.

Rajan, A., Tien, A. C., Haueter, C. M., Schulze, K. L. and Bellen, H. J. (2009). The Arp2/3 complex and WASp are required for apical trafficking of Delta into microvilli during cell fate specification of sensory organ precursors. Nature cell biology 11, 815–824.

Rakic, P. (2003a). Developmental and evolutionary adaptations of cortical radial glia. Cerebral cortex 13, 541–549.

Rakic, P.(2003b). Elusive radial glial cells: historical and evolutionary perspective. Glia 43, 19–32.

Rasin, M. R., Gazula, V. R., Breunig, J. J., Kwan, K. Y., Johnson, M. B., Liu-Chen, S., Li, H. S., Jan, L. Y., Jan, Y. N., Rakic, P., et al. (2007). Numb and Numbl are required for maintenance of cadherin-based adhesion and polarity of neural progenitors. Nature neuroscience 10, 819–827.

Robinson, R. C., Turbedsky, K., Kaiser, D. A., Marchand, J. B., Higgs, H. N., Choe, S. and Pollard, T. D. (2001). Crystal structure of Arp2/3 complex. Science 294, 1679–1684.

Rocca, D. L., Amici, M., Antoniou, A., Blanco Suarez, E., Halemani, N., Murk, K., McGarvey, J., Jaafari, N., Mellor, J. R., Collingridge, G. L., et al. (2013). The small GTPase Arf1 modulates Arp2/3-mediated actin polymerization via PICK1 to regulate synaptic plasticity. Neuron 79, 293–307.

Saedler, R., Mathur, N., Srinivas, B. P., Kernebeck, B., Hulskamp, M. and Mathur, J. (2004). Actin control over microtubules suggested by DISTORTED2 encoding the Arabidopsis ARPC2 subunit homolog. Plant & cell physiology 45, 813–822.

San Miguel-Ruiz, J. E. and Letourneau, P. C. (2014). The role of Arp2/3 in growth cone actin dynamics and guidance is substrate dependent. The Journal of neuroscience: the official journal of the Society for Neuroscience 34, 5895–5908.

Schmechel, D. E. and Rakic, P. (1979). A Golgi study of radial glial cells in developing monkey telencephalon: morphogenesis and transformation into astrocytes. Anatomy and embryology 156, 115–152.

Shakir, M. A., Jiang, K., Struckhoff, E. C., Demarco, R. S., Patel, F. B., Soto, M. C. and Lundquist, E. A. (2008). The Arp2/3 activators WAVE and WASP have distinct genetic interactions with Rac GTPases in Caenorhabditis elegans axon guidance. Genetics 179, 1957–1971.

Sheen, V. L. (2012). Periventricular Heterotopia: Shuttling of Proteins through Vesicles and Actin in Cortical Development and Disease. Scientifica 2012, 480129.

Shivas, J. M. and Skop, A. R. (2012). Arp2/3 mediates early endosome dynamics necessary for the maintenance of PAR asymmetry in Caenorhabditis elegans. Molecular biology of the cell 23, 1917–1927.

Spedden, E., White, J. D., Naumova, E. N., Kaplan, D. L. and Staii, C. (2012). Elasticity maps of living neurons measured by combined fluorescence and atomic force microscopy. Biophys J 103, 868–877.

Spillane, M., Ketschek, A., Jones, S. L., Korobova, F., Marsick, B., Lanier, L., Svitkina, T. and Gallo, G. (2011). The actin nucleating Arp2/3 complex contributes to the formation of axonal filopodia and branches through the regulation of actin patch precursors to filopodia. Developmental neurobiology 71, 747–758.

Strasser, G. A., Rahim, N. A., VanderWaal, K. E., Gertler, F. B. and Lanier, L. M. (2004). Arp2/3 is a negative regulator of growth cone translocation. Neuron 43, 81–94.

Suraneni, P., Fogelson, B., Rubinstein, B., Noguera, P., Volkmann, N., Hanein, D., Mogilner, A. and Li, R. (2015). A mechanism of leading-edge protrusion in the absence of Arp2/3 complex. Molecular biology of the cell 26, 901–912.

Suraneni, P., Rubinstein, B., Unruh, J. R., Durnin, M., Hanein, D. and Li, R. (2012). The Arp2/3 complex is required for lamellipodia extension and directional fibroblast cell migration. The Journal of cell biology 197, 239–251.

Tang, V. W. and Brieher, W. M. (2012). alpha-Actinin-4/FSGS1 is required for Arp2/3-dependent actin assembly at the adherens junction. The Journal of cell biology 196, 115–130.

Taylor, M. D., Poppleton, H., Fuller, C., Su, X., Liu, Y., Jensen, P., Magdaleno, S., Dalton, J., Calabrese, C., Board, J., et al. (2005). Radial glia cells are candidate stem cells of ependymoma. Cancer cell 8, 323–335.

Toret, C. P. and Drubin, D. G. (2006). The budding yeast endocytic pathway. Journal of cell science 119, 4585–4587.

Verma, S., Han, S. P., Michael, M., Gomez, G. A., Yang, Z., Teasdale, R. D., Ratheesh, A., Kovacs, E. M., Ali, R. G. and Yap, A. S. (2012). A WAVE2-Arp2/3 actin nucleator apparatus supports junctional tension at the epithelial zonula adherens. Molecular biology of the cell 23, 4601–4610.

Voss, A. K., Britto, J. M., Dixon, M. P., Sheikh, B. N., Collin, C., Tan, S. S. and Thomas, T. (2008). C3G regulates cortical neuron migration, preplate splitting and radial glial cell attachment. Development 135, 2139–2149.

Wegner, A. M., Nebhan, C. A., Hu, L., Majumdar, D., Meier, K. M., Weaver, A. M. and Webb, D. J. (2008). N-wasp and the arp2/3 complex are critical regulators of actin in the development of dendritic spines and synapses. The Journal of biological chemistry 283, 15912–15920.

Weitzdoerfer, R., Fountoulakis, M. and Lubec, G. (2002). Reduction of actin-related protein complex 2/3 in fetal Down syndrome brain. Biochemical and biophysical research communications 293, 836–841.

Wichterle, H., Garcia-Verdugo, J. M., Herrera, D. G. and Alvarez-Buylla, A. (1999). Young neurons from medial ganglionic eminence disperse in adult and embryonic brain. Nature neuroscience 2, 461–466.

Winter, D. C., Choe, E. Y. and Li, R. (1999). Genetic dissection of the budding yeast Arp2/3 complex: a comparison of the in vivo and structural roles of individual subunits. Proceedings of the National Academy of Sciences of the United States of America 96, 7288–7293.

Wu, C., Asokan, S. B., Berginski, M. E., Haynes, E. M., Sharpless, N. E., Griffith, J. D., Gomez, S. M. and Bear, J. E. (2012). Arp2/3 is critical for lamellipodia and response to extracellular matrix cues but is dispensable for chemotaxis. Cell 148, 973–987.

Yae, K., Keng, V. W., Koike, M., Yusa, K., Kouno, M., Uno, Y., Kondoh, G., Gotow, T., Uchiyama, Y., Horie, K., et al. (2006). Sleeping beauty transposon-based phenotypic analysis of mice: lack of Arpc3 results in defective trophoblast outgrowth. Molecular and cellular biology 26, 6185–6196.

Yang, Q., Zhang, X. F., Pollard, T. D. and Forscher, P. (2012). Arp2/3 complex-dependent actin networks constrain myosin II function in driving retrograde actin flow. The Journal of cell biology 197, 939–956.

Yokota, Y., Eom, T. Y., Stanco, A., Kim, W. Y., Rao, S., Snider, W. D. and Anton, E. S. (2010). Cdc42 and Gsk3 modulate the dynamics of radial glial growth, inter-radial glial interactions and polarity in the developing cerebral cortex. Development 137, 4101–4110.

Yokota, Y., Kim, W. Y., Chen, Y., Wang, X., Stanco, A., Komuro, Y., Snider, W. and Anton, E. S. (2009). The adenomatous polyposis coli protein is an essential regulator of radial glial polarity and construction of the cerebral cortex. Neuron 61, 42–56.

Zhang, J., Woodhead, G. J., Swaminathan, S. K., Noles, S. R., McQuinn, E. R., Pisarek, A. J., Stocker, A. M., Mutch, C. A., Funatsu, N. and Chenn, A. (2010). Cortical neural precursors inhibit their own differentiation via N-cadherin maintenance of beta-catenin signaling. Developmental cell 18, 472–479.

Zuo, X., Zhang, J., Zhang, Y., Hsu, S. C., Zhou, D. and Guo, W. (2006). Exo70 interacts with the Arp2/3 complex and regulates cell migration. Nature cell biology 8, 1383–1388.

